# Uncoupling of DNA replication and centrosome duplication cycles is a primary cause of haploid instability in mammalian somatic cells

**DOI:** 10.1101/2020.05.30.124826

**Authors:** Koya Yoshizawa, Kan Yaguchi, Ryota Uehara

## Abstract

Mammalian haploid somatic cells are unstable and prone to diploidize, but the cause of haploid instability remains largely unknown. Previously, we found that mammalian haploid somatic cells suffer chronic centrosome loss stemming from the uncoupling of DNA replication and centrosome duplication cycles. However, the lack of methodology to restore the coupling between DNA replication and centrosome duplication has precluded us from investigating the potential contribution of the haploidy-linked centrosome loss to haploid instability. In this study, we developed an experimental method that allows the re-coupling of DNA and centrosome cycles through the chronic extension of the G1/S phase without compromising cell proliferation using thymidine treatment/release cycles. Chronic extension of G1/S restored normal mitotic centrosome number and mitotic control, substantially improving the stability of the haploid state in HAP1 cells. Stabilization of the haploid state was compromised when cdk2 was inhibited during the extended G1/S, or when early G1 was chronically extended instead of G1/S, showing that the coupling of DNA and centrosome cycles rather than a general extension of the cell cycle is required for haploid stability. Our data indicate the chronic centriole loss arising from the uncoupling of centrosome and DNA cycles as a direct cause of genome instability in haploid somatic cells, and also demonstrate the feasibility of modulation of haploid stability through artificial coordination between DNA and centrosome cycles in mammalian somatic cells.

## Introduction

In animal organisms, haploid or near-haploid somatic cells arise from pathological processes such as parthenogenesis or aberrant chromosome loss in tumorigenesis (Wutz, 2014). The haploid state is unstable in mammalian somatic cells, and spontaneous whole-genome duplication converts haploid cells or embryos into diploid in several days or a few weeks (Kaufman et al., 1983; Andersson et al., 1995; Yang et al., 2013). This is in sharp contrast to plant or fungus cells, which in general proliferate stably in the haploid state. Though mammalian haploid cells are promising tools for mammalian genomics and engineering, their unstable nature limits the utility of haploid cell technology (He et al., 2019). The cause of haploid instability in mammalian somatic cells remains largely elusive. Several recent studies have found that haploid mammalian somatic cells commonly suffer severe mitotic delay and/or cell division failure, which potentially makes an important contribution to the progression of diploidization (Guo et al., 2017; He et al., 2017; Olbrich et al., 2017; Yaguchi et al., 2018). Interestingly, these mitotic defects are not seen or much less in their diploidized counterparts, hence haploidy-specific. However, the basis of these haploidy-specific defects remains obscure.

In mitosis, two centrosomes support bipolar spindle formation, which mediates equal segregation of sister chromatids. An abnormal number of centrosomes perturbs proper chromosome segregation, compromising genome integrity in daughter cells. To keep constant centrosome number over cell cycle generations, the progression of centrosome duplication is tightly coupled with that of DNA replication under control of cyclin E-cdk2 (Hinchcliffe et al., 1999; Lacey et al., 1999; Matsumoto et al., 1999; Meraldi et al., 1999; Fu et al., 2015). Previously, we found that centrosome duplication was drastically delayed in haploid HAP1 cells compared to their isogenic diploid counterparts, whereas the progression of DNA replication was almost equivalent between these ploidy states (Yaguchi et al., 2018). This uncoupling of DNA and centrosome cycles lead to chronic centrosome loss and frequent monopolar spindle formation specifically in haploid cells (Yaguchi et al., 2018), potentially affecting haploid stability. However, lack of experimental tools that enable the restoration of normal centrosome number and/or spindle polarity without compromising long-term cell viability has precluded us from directly testing causality of haploidy-associated centrosome loss for haploid instability (Yaguchi et al., 2018).

Here, we developed an experimental method to artificially recouple DNA and centrosome cycles by repeating the treatment and removal of thymidine. Using this method, we addressed whether the uncoupling of DNA and centrosome cycles in haploid cells is a primary cause of haploid stability in mammalian somatic cells.

## Method

### Cell culture and flow cytometry

Haploid and diploid HAP1 cells were cultured in Iscove’s Modified Dulbecco’s Medium (IMDM; Wako Pure Chemical Industries) supplemented with 10% fetal bovine serum and 1× antibiotic-antimycotic (Sigma-Aldrich). Haploid cells were regularly maintained by size-based cell sorting as previously described (Yaguchi et al., 2018). For DNA content analyses, cells were stained with 10 μg/ml Hoechst 33342 (Dojindo) for 15 min at 37°C, and fluorescence intensity was analyzed using a JSAN desktop cell sorter (Bay bioscience).

### Intermittent cell cycle blockage

In each inhibitor treatment/release cycle, cells were treated with 500 μM thymidine, 1 μM PD-0332991, or 1 μM LY-2835219 for 16 h, then rinsed with and incubated in supplemented IMDM without the inhibitors for 8 h. In the case of roscovitine co-treatment with thymidine, 5 μM roscovitine was introduced 5 h after the introduction of thymidine, treated for 11 h, and then washed out along with thymidine. Roscovitine was introduced from the second cycle of the thymidine treatment/release cycles. Cell passage was conducted using 0.05% trypsin-EDTA (Wako) typically once two days while cells were cultured in inhibitors-free medium. For immunofluorescence imaging, cells were fixed and stained 8 h after the release from the 3^rd^ inhibitor treatment. For live imaging, the cell culture medium was exchanged to supplemented phenol red-free IMDM (Thermo Fisher Scientific), and cells were imaged from 2 h after the release from the 3^rd^ inhibitor treatment. For monitoring the dynamics of DNA content in the long-term culture experiments, cells were subjected to flow cytometry 8 h after release from inhibitor treatment at the cycles of inhibitor treatment/release indicated in the main text and Fig. 3.

### Immunofluorescence (IF) staining

Cells were fixed with 100% methanol at −20°C for 10 min. Fixed samples were treated with blocking buffer (150 mM NaCl; 10 mM Tris-HCl, pH 7.5; 5% bovine serum albumin; and 0.1% Tween 20) for 30 min at 25°C, and incubated with primary and secondary antibodies overnight each at 4°C. Following each treatment, cells were washed 2–3 times with Dulbecco’s phosphate-buffered saline. Rat monoclonal anti-α-tubulin (YOL1/34, EMD Millipore); mouse monoclonal anti-centrin (20H5, EMD Millipore), goat anti-mouse Alexa Fluor 488-conjugated (ab150117, Abcam), and goat anti-rat Alexa Fluor 568-conjugated (ab175710, Abcam) antibodies were used at a dilution of 1:1000.

### Microscopy

Fixed cells were observed at 25°C under a TE2000 microscope (Nikon) equipped with a ×100 1.4 numerical aperture (NA) Plan-Apochromatic objective lens (Nikon), a CSU-X1 confocal unit (Yokogawa), and an iXon3 electron multiplier-charge coupled device camera (Andor). Living cells were observed for 24 h at 37°C with 5% CO_2_ under a Ti-2 microscope (Nikon) equipped with a ×40 0.95 NA Plan-Apochromatic objective lens (Nikon), and Zyla4.2 sCMOS camera (Andor). Sir-tubulin (Cytoskeleton, Inc.) was treated at 250 nM for live imaging. Image acquisition was controlled by µManager (Open Imaging).

### Colorimetric cell proliferation assay

For cell viability assay, 1,350 (haploid) or 675 (diploid) cells were seeded on each well of 96-well plates. After 24 h, cells were treated with different concentrations of thymidine. Forty-four h after the thymidine addition, 5% Cell Counting Kit-8 (Dojindo) was added to the culture, incubated for 4 h, and absorbance at 450 nm was measured using the Sunrise plate reader (Tecan). The absorbances of thymidine treated cells were normalized to those of the corresponding non-treated controls.

### Mathematical modeling and simulations

A mathematical cell population transition model was constructed as described previously with modifications (Yaguchi et al., 2018). The model is based on the following assumptions: 1) Haploid and diploid populations proliferate exponentially with characteristic doubling times corresponding to their cell cycle lengths; 2) in the case of intermittent cell cycle arrest, both haploid and diploid cells double once every 24 h; 3) haploid cells undergo mitotic death or convert into diploid through mitotic slippage with the observed frequencies; 4) G1 proportion is unchanged within haploid populations throughout the long-term culture. Assuming ergodicity in the processes of the cell cycle, mitosis, and cell proliferation, the probability of the occurrence of mitotic death or mitotic slippage in a haploid cell population during unit time length, *p*_*death*_ or *p*_*slippage*_, respectively, is derived as:

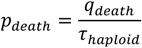

Or

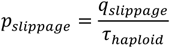

, respectively, where *q*_*slippage*_ and *q*_*death*_ are the rates of incidence of mitotic slippage and mitotic death per mitotic event, respectively, and τ_haploid_ is the average cell cycle length of a haploid cell population. The time-dependent growth of haploid or diploid cell population is modeled as:

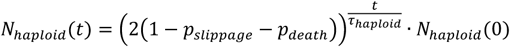

Or

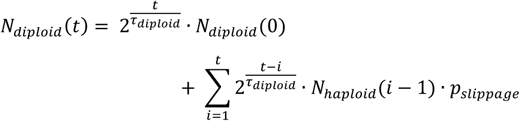

, respectively, where *N*_*haploid*_ or *N*_*diploid*_ is number of haploid or diploid cells, respectively, at a time point, *t*. τ_diploid_ is the average cell cycle length of a diploid cell population. A haploid cell that converts to diploid through mitotic slippage at any point during the simulation will thereafter proliferate with the characteristic doubling time of diploid cells. The percentage of haploid cells in G1, *P*_*haploid, G1*_, in cell culture is modeled as:

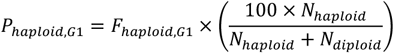

where *F*_*haploid, G1*_ is the fraction of cells in the G1 phase within a haploid cell population. Computer programs were written using MATLAB (Mathworks). Parameters used in simulations are listed in Table S1.

## Results

### Restoration of mitotic control by an artificial slowing of G1/S phase in haploid HAP1 cells

We reasoned that if the uncoupling of DNA replication and centrosome duplication cycles is a primary cause of mitotic defects and genome instability in haploid HAP1 cells, these haploidy-linked defects would be circumvented by artificial recoupling of DNA and centrosome cycles. Previously, we found that blockage of G1/S by thymidine allowed the continuous progression of centriole duplication in HAP1 (Yaguchi et al., 2018). Therefore, we first established an experimental method that chronically delays the DNA replication cycle and recouples it to the centrosome duplication cycle by thymidine. To extend G1/S in each round of cell cycle without halting cell proliferation, we intermittently repeated cycles of 16-h thymidine treatment followed by the subsequent 8-h incubation without thymidine (Fig. 1A; note that we previously quantified average cell cycle length of haploid HAP1 cells to be 13.4 h) (Yaguchi et al., 2018). Using flow cytometry, we confirmed that cell cycle progression was effectively blocked and released by thymidine treatment/removal cycles (Fig. S1A and B). Thymidine cycles resulted in a significant increase in cell size compared to mock-treated control, while the cells kept the haploid DNA content (Fig. S1C). Next, we tested the effects of thymidine cycles on mitotic spindle organization and mitotic progression in the haploid state by live imaging of HAP1 cells stained with a fluorescent microtubule marker SiR-tubulin (Fig. 1B). Most of the diploid control cells formed bipolar spindle by 20 min after nuclear envelope breakdown (NEBD) and entered anaphase by 35 min after NEBD (Fig. 1C and D). In contrast, a substantial proportion of haploid mitotic cells showed a severe delay in or absence of monopolar-to-bipolar conversion, leading to drastic mitotic arrest (Fig. 1C and D). Out of 24 haploid cells whose mitotic consequences could be specified after drastic mitotic arrest (>50 min), 3 or 5 cells underwent mitotic death or mitotic slippage (mitotic exit without cytokinesis), respectively (Fig. 1E). In contrast, these mitotic defects were not observed in diploid control (Fig. 1E). Interestingly, after three cycles of thymidine treatment/release in haploid cells, the frequency of the delay and/or blockage of monopolar-to-bipolar conversion was drastically reduced (Fig. 1C). Thymidine cycles also restored normal mitotic progression with substantial suppression of mitotic slippage (Fig. 1D and E). To investigate the effects of thymidine cycles on mitotic centrosome number, we also conducted immunostaining of centrin-2 (centriole marker) and α-tubulin (spindle marker) in non-treated or thymidine cycles-treated haploid cells (Fig. 2A-D). Whereas non-treated haploid cells showed frequent centriole loss accompanied by spindle monopolarization, both normal mitotic centriole number and spindle bipolarity were significantly restored after three cycles of thymidine treatment/release (Fig. 2C and D). Therefore, artificial slowing of G1/S recoupled DNA and centrosome cycles, largely improving mitotic control in haploid cells.

**Figure 1:**
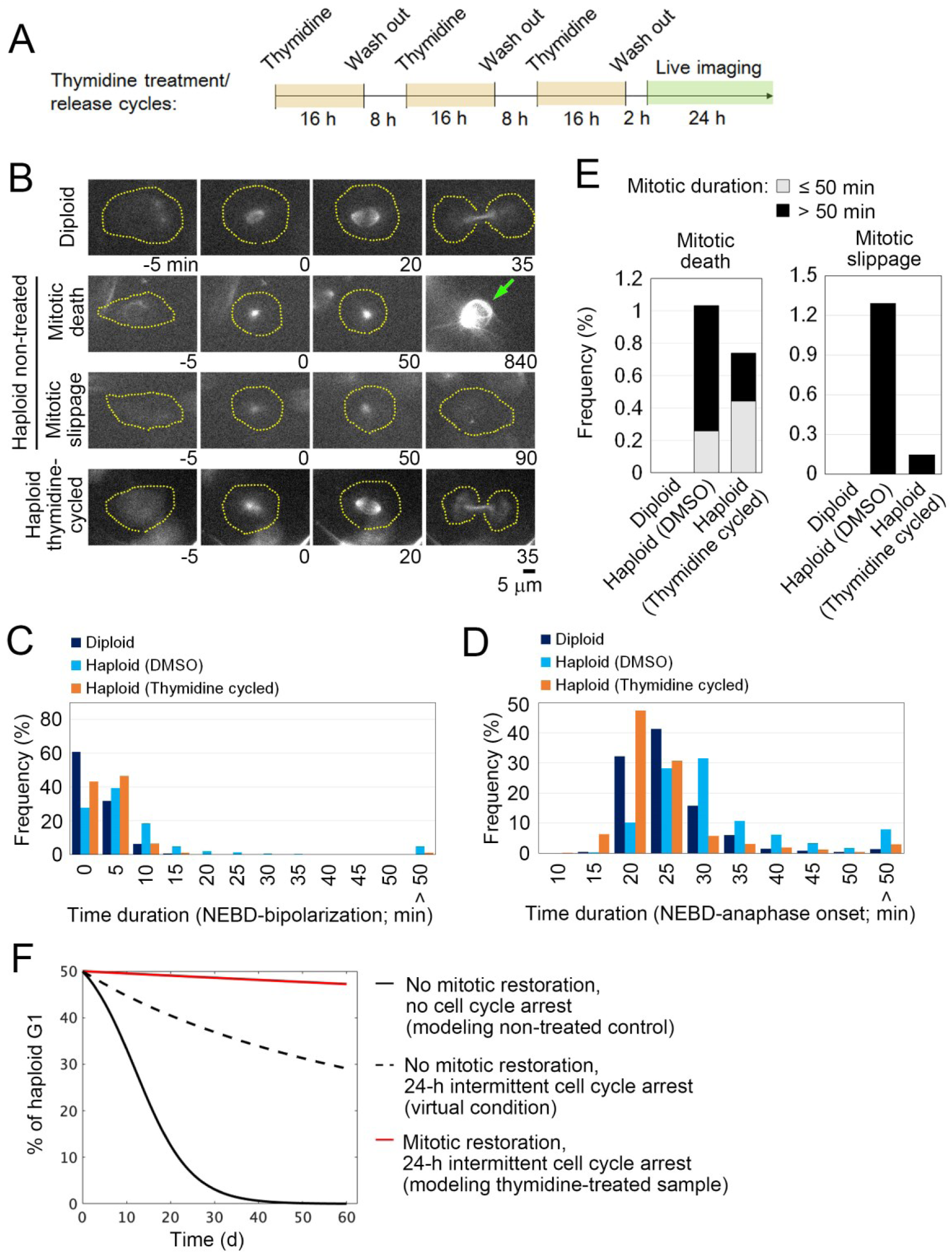
Restoration of mitotic progression by intermittent extension of G1/S phase in haploid HAP1 cells. (**A**) Experimental scheme of thymidine treatment/release cycles. (**B**) Live images of SiR-tubulin-stained HAP1. Images were taken at 5-min intervals. NEBD was set as 0 min. Broken lines: cell boundaries. Arrow: a dead cell. (**C, D, E**) Distribution of time duration for monopolar-to-bipolar conversion (C) or NEBD-to-anaphase transition (D), or the frequency of mitotic defects (E). At least 393 (C, D) or 387 cells (E) from two independent experiments were analyzed. (**F**) Theoretical model simulation of haploid instability in long-term culture with or without intermittent cell cycle arrest and/or mitotic restoration.

**Figure 2:**
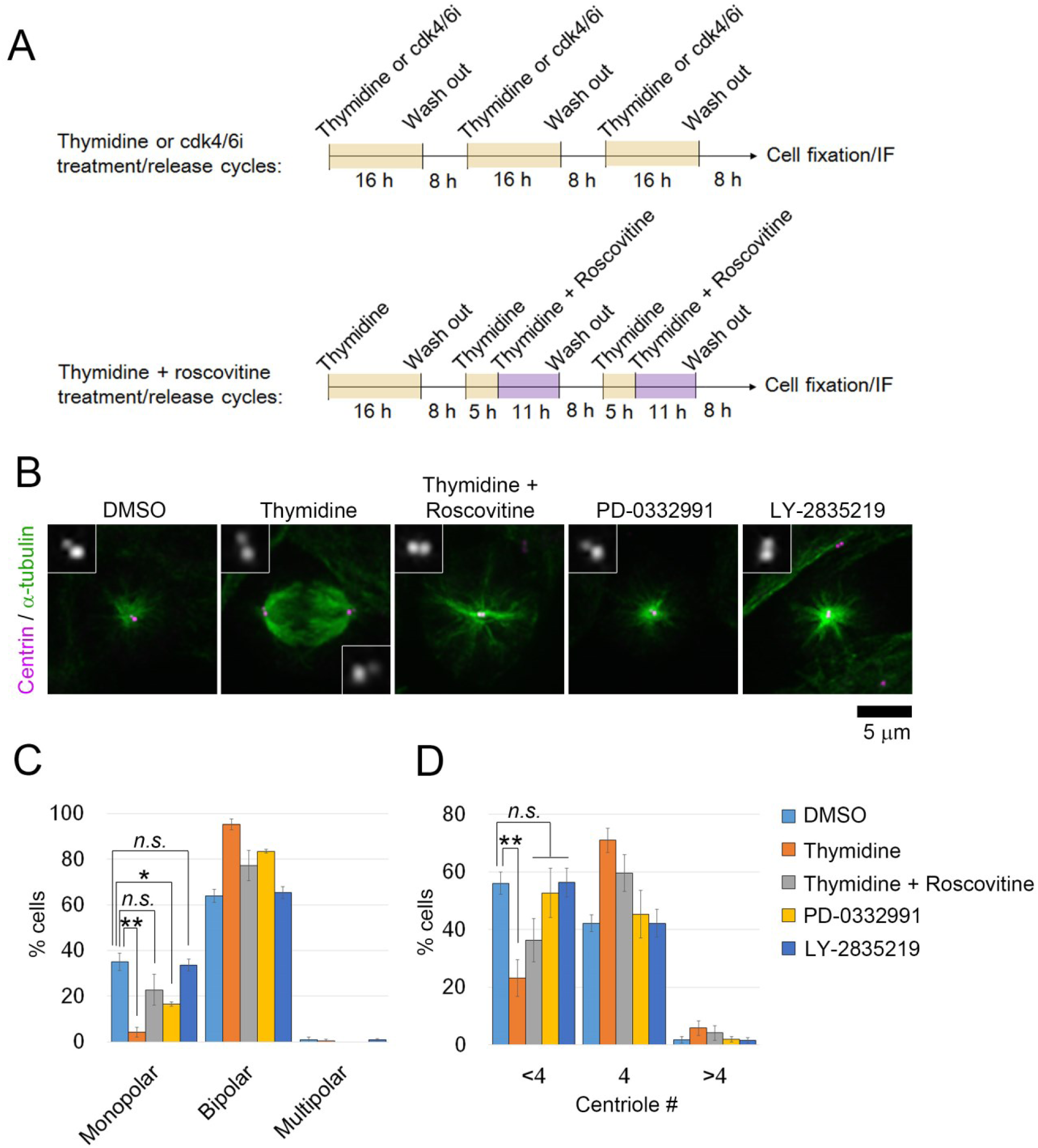
Restoration of normal mitotic centrosome number and spindle bipolarity by intermittent extension of G1/S phase in haploid HAP1 cells. (**A**) Experimental scheme of inhibitors treatment/release cycles. (**B**) Immunostaining of α-tubulin and centrin in haploid cells after three cycles of inhibitors treatment/release. Insets show 3× enlarged images of the centrioles. (**C, D**) Frequency of spindle polarities (C) or centriole numbers (D). Means ± standard error (SE) of three independent experiments (* p < 0.05, ** p < 0.01, two-tailed *t*-test). At least 154 cells were analyzed per condition.

### Improvement of haploid stability in long-term culture by the recoupling of DNA and centrosome cycles

We next theoretically estimated impacts of the thymidine-mediated improvement of mitotic control on haploid stability using a mathematical cell population model developed in our previous study (Yaguchi et al., 2018) (*Method*). The model simply assumes that haploid and diploid cells proliferate exponentially with their characteristic doubling times, while haploid cells undergo mitotic death or diploidize through mitotic slippage at the frequencies quantified in the live imaging (Fig. 1E). We previously found that diploid HAP1 cells proliferate faster than haploid cells with a cell cycle length of 11.9 h (vs 13.4 h in haploid), which promotes the expansion of diploidized cells over haploid (Yaguchi et al., 2018). To test the potential influence of intermittent cell cycle arrest on the haploid-to-diploid conversion, we first compared a theoretical condition in which haploid and diploid cells proliferate with their observed doubling times with another condition in which both of them double only once every 24 h (Fig. 1F). Even when we assumed the identical frequencies of mitotic death and mitotic slippage in these two conditions, the haploid-to-diploid conversion was much slowed down by intermittent cell cycle arrest (Fig. 1F and Table S1). Therefore, the reduction of mitotic frequency and the cancellation of the ploidy-dependent growth difference by the intermittent cell cycle arrest *per se* make a substantial contribution to haploid stability. Then, we compared the haploid-to-diploid conversion between theoretical conditions in which mitotic death and mitotic slippage occurred at different frequencies as observed between thymidine cycles-treated and non-treated haploid cells. The simulation showed that the observed level of mitotic restoration by thymidine cycles could substantially improve haploid stability even if we removed from consideration the effect of intermittent cell cycle arrest (compare broken black and red lines in Fig. 1E). Therefore, the modulation of mitotic control through the recoupling of DNA and centrosome cycles was predicted to profoundly improve the long-term stability of the haploid state.

Next, we experimentally tested the model prediction. To evaluate the net contribution of mitotic restoration to haploid stability independently from that of intermittent cell cycle arrest, we needed to set control conditions in which cell cycle was intermittently arrested whereas DNA and centrosome cycles remained uncoupled. For this, i) a cdk2 inhibitor roscovitine was co-treated in thymidine cycles to inhibit cdk2-mediated centriole duplication in G1/S (Hinchcliffe et al., 1999; Lacey et al., 1999; Matsumoto et al., 1999; Meraldi et al., 1999), or ii) early G1 phase was intermittently blocked instead of G1/S by cdk4/6 inhibitor (PD-0332991 or LY-2835219) treatment/release cycles (Fry et al., 2004; Gelbert et al., 2014) (Fig. 2A). In the case of thymidine and roscovitine co-treatment, we had to avoid blockage of mitotic progression by roscovitine-mediated cdk1 inhibition. For this, roscovitine was introduced at 13 h after the previous release from thymidine arrest, which was sufficient for the majority of cells to pass through the mitotic phase and to be arrested at the next G1/S boundary (Fig. S1A and B). Cell cycle profile and cell size distribution were almost identical between cells in the thymidine cycles and those in thymidine and roscovitine cycles (Fig. S1B and C). However, roscovitine substantially attenuated the restoration of mitotic centriole number and spindle bipolarity in thymidine cycles-treated haploid cells (Fig. 2B-D). Similarly, though PD-0332991 or LY-2835219 cycles effectively blocked and released cell cycle progression (Fig. S1B), restoration of mitotic centrosome number and spindle bipolarity in the cells treated with these inhibitors was scarce or ignorable (Fig. 2B-D). Therefore, restoration of the coupling between DNA replication and centrosome duplication was specifically achieved by the G1/S phase extension in a cdk2-dependent manner. These results demonstrate that the above experimental conditions serve as controls that intermittently block cell cycle progression while keeping DNA and centrosome cycles uncoupled in haploid cells.

We next conducted long-term continuous passages while intermittently blocking early G1 or G1/S by cdk4/6 inhibitors or thymidine, respectively, on the same schedule of the cycle as described in Fig. 2A. After continuous passages for 30 d, non-treated control haploid cells drastically diploidized, which was marked by a substantial reduction in haploid G1 population (indicated as “1C” peak in flow cytometry in Fig. 3A) along with the emergence of diploid G2/M population (“4C” peak); the proportion of haploid G1 reduced to 5.9% on day 30 (Fig. 3B). Thymidine cycles drastically suppressed diploidization with the proportion of haploid G1 remaining to be 37% on day 30 (Fig. 3A and B). The suppression of diploidization was much milder or ignorable in cdk4/6 inhibitors-treated cells, excluding the possibility that haploid stability was improved merely because of delayed cell proliferation in thymidine cycles. Moreover, co-treatment of roscovitine substantially attenuated the thymidine-mediated haploid stabilization with the proportion of haploid G1 being 27% on day 30. These data support the idea that the uncoupling of DNA and centrosome cycles is a primary cause of haploid instability in HAP1. During the prolonged culture over 30 d with thymidine cycles, cells became insensitive to thymidine-mediated cell cycle arrest (Fig. 3C) (Morrow and Stocco, 1980), which precluded further preservation of the haploid cell population.

**Figure 3:**
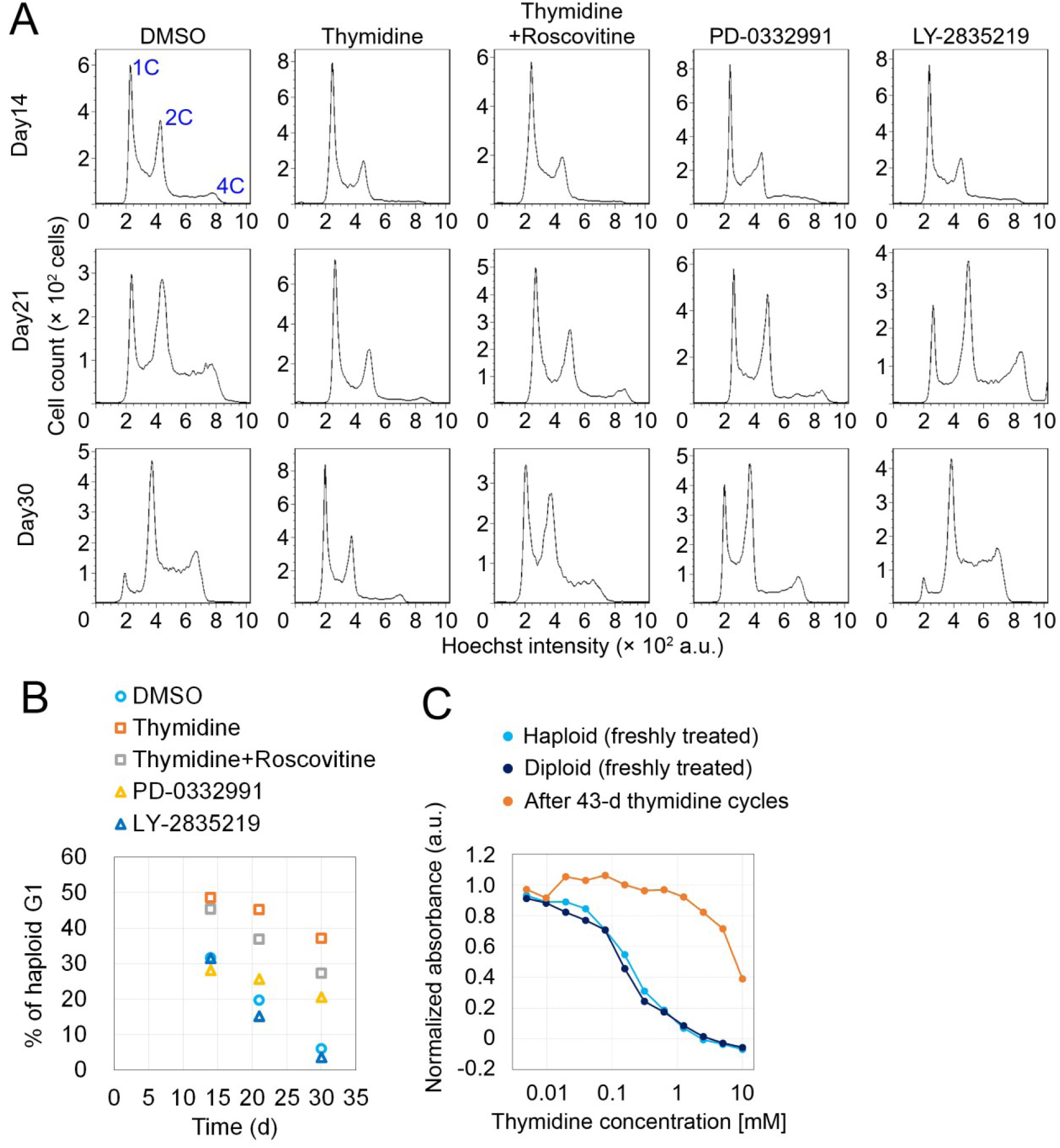
Improvement of haploidy stability in G1/S phase-extended cells in long-term passages. (**A**) Flow cytometric analysis of DNA content in Hoechst-stained cells in the long-term passages with or without inhibitors treatment/release cycles. Representative data from two independent experiments. (**B**) Time course of haploid G1 frequency in A. Means of two independent experiments. (**C**) Colorimetric cell proliferation assay of HAP1 cells treated with different concentrations of thymidine before or after long-term thymidine treatment/release cycles. A representative result from two independent experiments is shown (means of duplicates).

## Discussion

In this study, by chronically delaying G1/S progression by the intermittent thymidine treatment, we could establish, for the first time, an experimental condition in which mitotic centrosome number and mitotic control were substantially restored in human haploid somatic cells. Mitotic restoration drastically improved haploid stability. On the other hand, suppression of centrosome restoration by cdk2 inhibition attenuated the effect of G1/S extension on haploid stability. Our results indicate the causality between the haploidy-linked centrosome loss and the instability of the haploid state in human somatic cells.

Because of the rich potential of mammalian haploid cells in bioengineering, many attempts have been made to improve haploid stability. Different studies have reported that diploidization in mouse haploid ES cells is substantially suppressed by optimized inhibition of wee1 kinase, cdk1, Rho-kinase or GSK-3/TGF-β pathways/BMP4 pathway (Takahashi et al., 2014; He et al., 2017; Li et al., 2017), or by gene manipulations including ectopic expression of aurora B kinase or Dnmt3b (Guo et al., 2017; He et al., 2018). Intriguingly, the effects of these treatments on haploid stability have been attributed to the acceleration of G2/M. Our results are consistent with this proposal in that we also observed a correlation between the restoration of normal mitotic progression and haploid stability. It was not determined how the acceleration of G2/M improved haploid stability in the previous studies. However, since our results suggest that the haploidy-specific mitotic slippage makes an important contribution to haploid instability, a possible explanation would be that G2/M acceleration stabilizes the haploid state by reducing the chance of mitotic slippage. Another possible mechanism of the G2/M acceleration-mediated haploid stabilization may be through the modulation of p53 state in haploid cells. A previous study reported haploidy-linked p53 upregulation, which limits the proliferation of haploid cells (Olbrich et al., 2017). Interestingly, recent studies have shown that prolonged mitosis arising from centrosome loss is sufficient to cause p53-dependent cell growth arrest (Fong et al., 2016; Lambrus et al., 2016; Meitinger et al., 2016). Therefore, we attempted to address whether centrosome restoration by thymidine cycles could suppress p53 upregulation in haploid cells. However, thymidine treatment *per se* triggered p53 upregulation presumably because of the replication folk stress (not shown) (Bolderson et al., 2004), precluding us from testing this idea.

Our study demonstrated the feasibility of haploid stabilization by the recoupling of DNA and centrosome cycles. However, to improve the versatility in routine cell maintenance, simpler methods for chronic recoupling of DNA and centrosome cycles are desirable. One possible approach would be to establish haploid mutant cell lines in which DNA replication or centrosome duplication is chronically delayed or accelerated, respectively, while cell viability remains unaffected. The loss-of-function of non-essential activators of DNA replication or promotion of centrosome duplication factors would be potential gene manipulation approaches. Our study provides a basis for an understanding of the mechanism that determines ploidy dynamics in mammalian somatic cells, as well as for further improvements of haploid cell technologies.

## Supporting information

modeling

## Acknowledgment

We thank Kimino Sato and Takahiro Yamamoto for technical assistance, and the Global Facility Center at Hokkaido University for the flow cytometer. This work was supported by Grant-in-Aid for JSPS Fellows to K.Yaguchi (19J12210), Grants-in-Aid for Scientific Research B (19H03219), on Innovative Areas “Singularity Biology (No.8007)” (19H05413), Fostering Joint International Research B (19KK0181), and JSPS Bilateral Joint Research Project (JPJSBP120193801) of MEXT, the Princess Takamatsu Cancer Research Fund, the Kato Memorial Bioscience Foundation, the Orange Foundation, the Smoking Research Foundation, the Suhara Memorial Foundation, and the Nakatani Foundation to R.Uehara.

## Author Contributions

Conceptualization, R.U.; Methodology, K.Yo., K.Ya., and R.U.; Investigation, K.Yo, K.Ya, and R.U.; Formal Analysis, K.Yo., K.Ya., and R.U.; Resources, R.U.; Writing – Original Draft, R.U.; Writing – Review & Editing, K. Yo., K. Ya., and R.U.; Funding Acquisition, K.Ya., and R.U.

---

**Figure S1:**
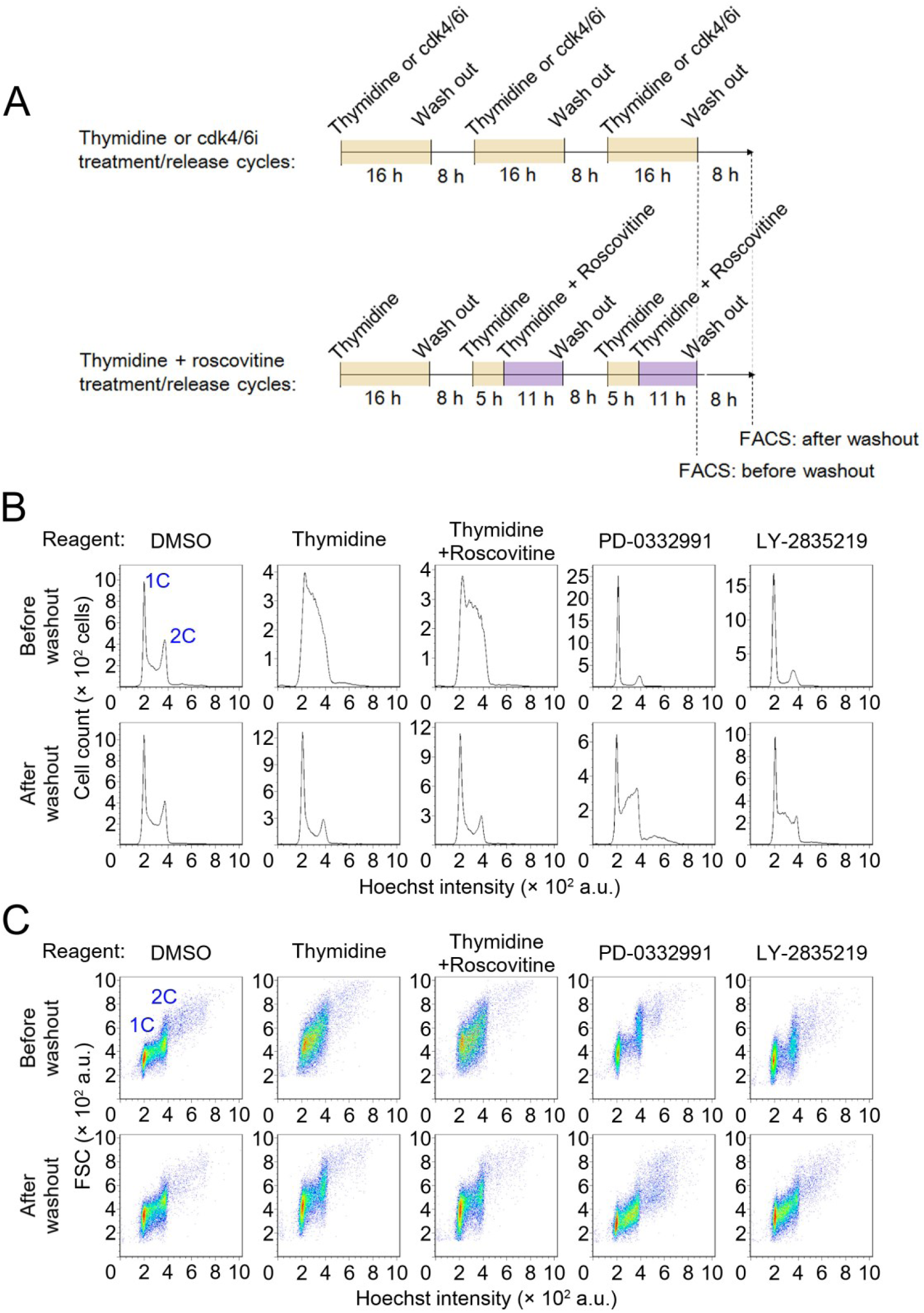
Intermittent extension of DNA replication cycle by cell cycle inhibitor treatment/release. (**A**) Experimental scheme of inhibitor treatment/release cycles. (**B, C**) Flow cytometric analysis of DNA content in Hoechst-stained haploid cells in inhibitors treatment/release cycles. Histograms of Hoechst signal and dot plots of forward scatter signal (for the judgment of relative cell size) against the Hoechst signal are shown in B and C, respectively. Representative data from three independent experiments. Note that co-treatment of roscovitine did not change the profile of cell cycle arrest/release in the thymidine treatment/release cycle.

**Table S1:** Parameters and constants used in the simulations

| Constants<br>(Unit) | Meaning | No mitotic<br>restoration,<br>no cell cycle arrest<br>(untreated<br>control) | No mitotic rescue,<br>24-h intermittent<br>cell cycle arrest<br>(Virtual condition) | Mitotic restoration,<br>24-h intermittent cell<br>cycle arrest<br>(thymidine-treated<br>sample) |
| --- | --- | --- | --- | --- |
| $N_{haploid}(0)$<br>($\times 10^3$ cells) | The initial<br>number of<br>haploid<br>cells | 5 | 5 | 5 |
| $N_{diploid}(0)$<br>($\times 10^3$ cells) | The initial<br>number of<br>diploid<br>cells | 0 | 0 | 0 |
| $q_{slippage}$<br>(h) | Incidence<br>rate of<br>mitotic<br>slippage<br>per mitotic<br>event | 0.012 | 0.012 | 0.001 |
| $q_{death}$<br>(h) | Incidence<br>rate of<br>mitotic<br>death per<br>mitotic<br>event | 0.010 | 0.010 | 0.007 |
| $\tau_{haploid}$<br>(h) | Cell cycle<br>length of<br>haploid | 13.4 | 24 | 24 |
| $\tau_{diploid}$<br>(h) | Cell cycle<br>length of<br>diploid | 11.9 | 24 | 24 |
| $F_{haploid, G1}$<br>(%) | G1 cell<br>fraction<br>within<br>haploid<br>population | 50 | 50 | 50 |

