## Supplementary material for "Uncoupling of DNA replication and centrosome duplication cycles is a primary cause of haploid instability in mammalian somatic cells": modeling: haploid.m

```
% calculate population
%

function [Hap Dip Hperc HapG1perc] = haploid(t)
H_cy = 24;
D_cy = 24;
Hap = 5000;
Dip = 0;
m = 0.001/H_cy;
c = 0.007/H_cy;
HapG1ratio =50;

% calculate population of each timing 
Hap_t(1) = Hap;
for i = 1:t
  Hap_t(i+1) = ((2*(1-m-c))^(i/H_cy))*Hap;  
  Dip_t(i) = (2^((t-i)/D_cy))*Hap_t(i)*m;
end

% calculate population at t
Hap = ((2*(1-m-c))^(t/H_cy))*Hap;
Dip = (2^(t/D_cy))*Dip + sum(Dip_t);
Hperc = Hap/(Dip+Hap) *100;
HapG1perc = HapG1ratio/100 * Hperc;
```
