## Supplementary material for "Uncoupling of DNA replication and centrosome duplication cycles is a primary cause of haploid instability in mammalian somatic cells": modeling: plot_hap.m

```
% plot population
%

function plot_hap(n)
m = zeros(n);
for i = 1:n
  [Hap(i) Dip(i) Hperc(i) HapG1perc(i)] = haploid(i);
  

  
end

figure;
%subplot(4,1,1);plot(Hap);title('Haploid #');
%subplot(4,1,2);plot(Dip);title('Diploid #');
%subplot(4,1,3);plot(Hperc);title('Hap total percntage');
%subplot(4,1,4);
plot(HapG1perc);title('HapG1 percentage');
```
